# Molecular Determinants of Complexin Clamping in Reconstituted Single-Vesicle Fusion

**DOI:** 10.1101/2021.07.05.451112

**Authors:** Manindra Bera, Sathish Ramakrishnan, Jeff Coleman, Shyam S. Krishnakumar, James E. Rothman

**Affiliations:** Departments of Cell Biology, Yale University School of Medicine, 333 Cedar Street, New Haven, CT 06520, USA; Department of Pathology, Yale University School of Medicine, 333 Cedar Street, New Haven, CT 06520, USA; Department of Neurology, Yale University School of Medicine, 333 Cedar Street, New Haven, CT 06520, USA

## Abstract

Previously we reported that Synaptotagmin-1 and Complexin synergistically clamp the SNARE assembly process to generate and maintain a pool of docked vesicles that fuse rapidly and synchronously upon Ca^2+^ influx (Ramakrishnan et al. 2020). Here using the same *in vitro* single-vesicle fusion assay, we establish the molecular details of the Complexin clamp and its physiological relevance. We find that a delay in fusion kinetics, likely imparted by Synaptotagmin-1, is needed for Complexin to block fusion. Systematic truncation/mutational analyses reveal that continuous alpha-helical accessory-central domains of Complexin are essential for its inhibitory function and specific interaction of the accessory helix with the SNAREpins, analogous to the *trans* clamping model, enhances this functionality. The c-terminal domain promotes clamping by locally elevating Complexin concentration through interactions with the membrane. Further, we find that Complexin likely contributes to rapid Ca^2+^-synchronized vesicular release by preventing un-initiated fusion rather than by directly facilitating vesicle fusion.

---

Neurons communicate with each other at synaptic contacts by releasing neurotransmitters from synaptic vesicles (SVs). This process is tightly controlled by activity-dependent changes in the presynaptic Ca^2+^ concentration and can occur in less than a millisecond after the neuronal spike (Sudhof 2013; Kaeser and Regehr 2014). SV fusion is catalyzed by presynaptic SNARE proteins. The SNAREs on the opposing membranes (VAMP2 on the synaptic vesicle membrane; Syntaxin and SNAP25 on the presynaptic plasma membrane) assemble into a four-helix bundle that catalyzes fusion by forcing the two membranes together (Sollner et al. 1993; Weber et al. 1998). Related SNARE proteins are universally involved in intracellular transport pathways and by themselves can constitutively catalyze fusion (Weber et al. 1998; McNew et al. 2000). As such, Ca^2+^-evoked neurotransmitter release occurs from the readily releasable pool (RRP) of vesicles docked/primed at the presynaptic active zone (Sudhof 2013; Kaeser and Regehr 2014). The current view is that at a single RRP vesicle, the SNARE complexes are firmly held (‘clamped’) in a partially assembled state (SNAREpins) close to the point of triggering fusion. Upon Ca^2+^ influx, multiple SNAREpins are synchronously activated to drive ultra-fast SV fusion and neurotransmitter release (Sudhof and Rothman 2009; Sudhof 2013; Rizo and Xu 2015; Rothman et al. 2017; Brunger et al. 2019).

It is well-established that the late stages of SV fusion are tightly regulated by two synaptic proteins – the presynaptic Ca^2+^ release sensor Synaptotagmin-1 (Syt1) and Complexin (CPX) (Sudhof 2013; Sudhof and Rothman 2009; Rizo and Xu 2015; Brunger et al. 2019). CPX is an evolutionarily conserved cytosolic protein that bind and regulate synaptic SNARE complex assembly (McMahon et al. 1995; Huntwork and Littleton 2007; Martin et al. 2011; Trimbuch and Rosenmund 2016; Mohrmann et al. 2015). Biochemical and biophysical analyses show that CPX promotes the initial stages of SNARE assembly but then blocks complete assembly (Li et al. 2011; Kummel et al. 2011; Lai et al. 2014; Krishnakumar et al. 2015). Thus, it can both facilitate and subsequently inhibit SV fusion. CPX contain distinct domains that mediate the dual clamp/activator function (Xue et al. 2007; Giraudo et al. 2008; Trimbuch and Rosenmund 2016; Mohrmann et al. 2015). The largely unstructured N-terminal domain (residues 1-26 of mammalian CPX1) activates Ca^2+^-regulated vesicular release (Xue et al. 2010; Lai et al. 2016) while the α-helical accessory domain (CPX_acc_, residues 26-48) serves as the primary clamping domain (Xue et al. 2007; Giraudo et al. 2008; Maximov et al. 2009; Yang et al. 2010; Kummel et al. 2011; Cho et al. 2014). A central helical sequence within CPX (CPX_cen_, residues 48-70) binds the groove between pre-assembled Syntaxin and VAMP2 and is essential for both function (Chen et al. 2002; Xue et al. 2007; Giraudo et al. 2008; Maximov et al. 2009). The remainder c-terminal portion (residues 71-134) has been shown to preferentially associate with curved lipid membrane via an amphipathic helical region and promotes the clamping function (Kaeser-Woo et al. 2012; Wragg et al. 2013; Gong et al. 2016).

The relative strength of CPX facilitatory vs inhibitory activities differs across species (Yang et al. 2013; Trimbuch and Rosenmund 2016; Mohrmann et al. 2015; Xue et al. 2009). As a result of this intricate balance, genetic perturbations of CPX can produce apparently contradictory effects in different systems. For example, knockout (KO) of CPX in neuromuscular synapses of *C. elegans* and *Drosophila* results in increased spontaneous release, decreased evoked release with overall reduction in the RPP size (Huntwork and Littleton 2007; Cho et al. 2014; Martin et al. 2011; Hobson et al. 2011; Wragg et al. 2013). In model mammalian synapses, CPX KO abates both spontaneous and evoked release with no significant change in the RRP size (Reim et al. 2001; Xue et al. 2008; Lopez-Murcia et al. 2019) but acute CPX knockdown (KD) reduces synaptic strength, but also increases spontaneous release with a concomitant reduction in the number of primed vesicles (Maximov et al. 2009; Yang et al. 2010; Kaeser-Woo et al. 2012; Yang et al. 2013). Some of the apparent discrepancies might be related to the perturbation method used (Yang et al. 2013), nonetheless, the physiological role of CPX in regulating SV fusion and the underlying mechanisms remains in the center of debate (Mohrmann et al. 2015; Trimbuch and Rosenmund 2016).

The interpretation of the physiological experiments can be limited by presence of the different CPX isoforms and possible compensatory homeostatic mechanisms. As such, the experiments in live synapses need to be complemented with a reductionist approach where the variables are limited, and the components can be rigorously controlled or altered. It is our hypothesis that the most direct mechanistic insight can be obtained from fully controlled cell-free systems. We have described a biochemically defined fusion setup based on a pore-spanning lipid bilayer setup that is best-suited for this purpose (Ramakrishnan et al. 2018; Ramakrishnan et al. 2019; Ramakrishnan et al. 2020).

Using this *in vitro* setup, which allows for precision study of the single-vesicle fusion kinetics, we recently demonstrated that mammalian CPX (mCPX), along with Syt1 and SNAREs, are essential and sufficient to achieve Ca^2+^-regulated fusion under physiologically-relevant conditions (Ramakrishnan et al. 2020). Our data revealed that mCPX and Syt1 act co-operatively to clamp the SNARE assembly process and produce a pool of docked vesicles. However, the study also revealed that there are at least two types of clamped SNAREpins under a docked vesicle – a small subset that are reversibly clamped by binding to Syt1 (which we termed ‘central’) and a larger population that are thought to be free of Syt1 and require mCPX for clamping (termed ‘peripheral’). We further established that Syt1s’ ability to oligomerize and bind SNAREpins via the ‘primary’ binding site on SNAP25 is key to its ability to clamp central SNAREpins and that the activation of these Syt1-associated SNAREpins is sufficient to elicit rapid, Ca^2+^-synchronized vesicle fusion(Ramakrishnan et al. 2020).

Building on this work, here we use a systematic *in vitro* reconstitution strategy to obtain new and direct insights into the molecular basis of mCPX clamping function and its role in establishing Ca^2+^-regulated release. We report that mCPX inhibitory function requires a delay in overall fusion kinetics and involves well-defined interaction of the accessory-central helical fragments with the SNAREpins. We further find that mCPX is essential to generate and maintain a pool of docked vesicles but does not contribute to the Ca^2+^-triggered rapid (<10 ms) and synchronous fusion of the docked vesicles.

## RESULTS

To dissect the mCPX clamping functionality, we used physiologically-relevant reconstitution conditions similar to our previous work (Ramakrishnan et al. 2020). Typically, we used small unilamellar vesicles (SUV) with in an average of 74 copies (outward facing) of VAMP2 (vSUV) without or with 25 copies Syt1 (Syt1-vSUV) (Figure 1 Supplement 1). We employed pre-formed t-SNAREs (1:1 complex of Syntaxin1 and SNAP-25) in the planar bilayers (containing 15% PS and 3% PIP2) to both simplify the experimental approach and to bypass the requirement of SNARE-assembling chaperones, Munc18 and Munc13 (Baker and Hughson 2016). Mammalian CPX1 (wild type or variants) was included in solution, typically at 2 μM unless noted otherwise. We used fluorescently labelled lipid (2% ATTO647N-PE) to track docking, clamping and spontaneous fusion of individual vesicles and a content dye (sulforhodamine B) to study Ca^2+^-triggered fusion of docked vesicles from the clamped state.

To focus on the ‘clamping’ of constitutive fusion events, we monitored large ensembles of vesicles to determine the percent remaining unfused as a function of time elapsed after docking and quantified as ‘survival percentages’ (Ramakrishnan et al. 2019; Ramakrishnan et al. 2020; Ramakrishnan et al. 2018). Docked immobile vesicles that remained un-fused during the initial 10 min observation period were defined as ‘clamped’ and the ‘docking-to-fusion’ delay enabled us to quantify the strength of the fusion clamp (Ramakrishnan et al. 2019; Ramakrishnan et al. 2020; Ramakrishnan et al. 2018). Since we track the fate of single vesicles, this analysis allowed us to examine the ‘clamping’ mechanism, independent of any alteration in the preceding docking sub-step.

Our earlier results showed that Syt1 without mCPX can meaningfully delay but not stably clamp SNARE-mediated fusion. Similarly, mCPX, on its own, is ineffective in clamping SNARE-driven vesicle fusion. In fact, both Syt1 and mCPX are needed to produce a stably ‘clamped’ state which can then be reversed by Ca^2+^ (Ramakrishnan et al. 2020). It is possible that Syt1 and mCPX1 either act jointly to generate a new intermediate state in the SNARE assembly pathway or operate sequentially, with the kinetic delay introduced by Syt1 enabling mCPX to arrest SNARE assembly. To distinguish between these possibilities, we developed a mimic for the Syt1 clamp – a lipid-conjugated ssDNA that is capable of regulating SNARE-driven fusion *in situ*. Without directly interacting with the SNAREs, the specific base-pair hybridization of the complementary ssDNA reconstituted into the SUVs and the planar bilayer introduces a steric barrier which is expected to, and indeed does delay fusion (Figure 1, Figure 1 Supplement 2). Moreover, this docking-to-fusion delay could be varied by adjusting the number of ssDNA molecules (Figure 1 Supplement 2).

**Figure 1.**
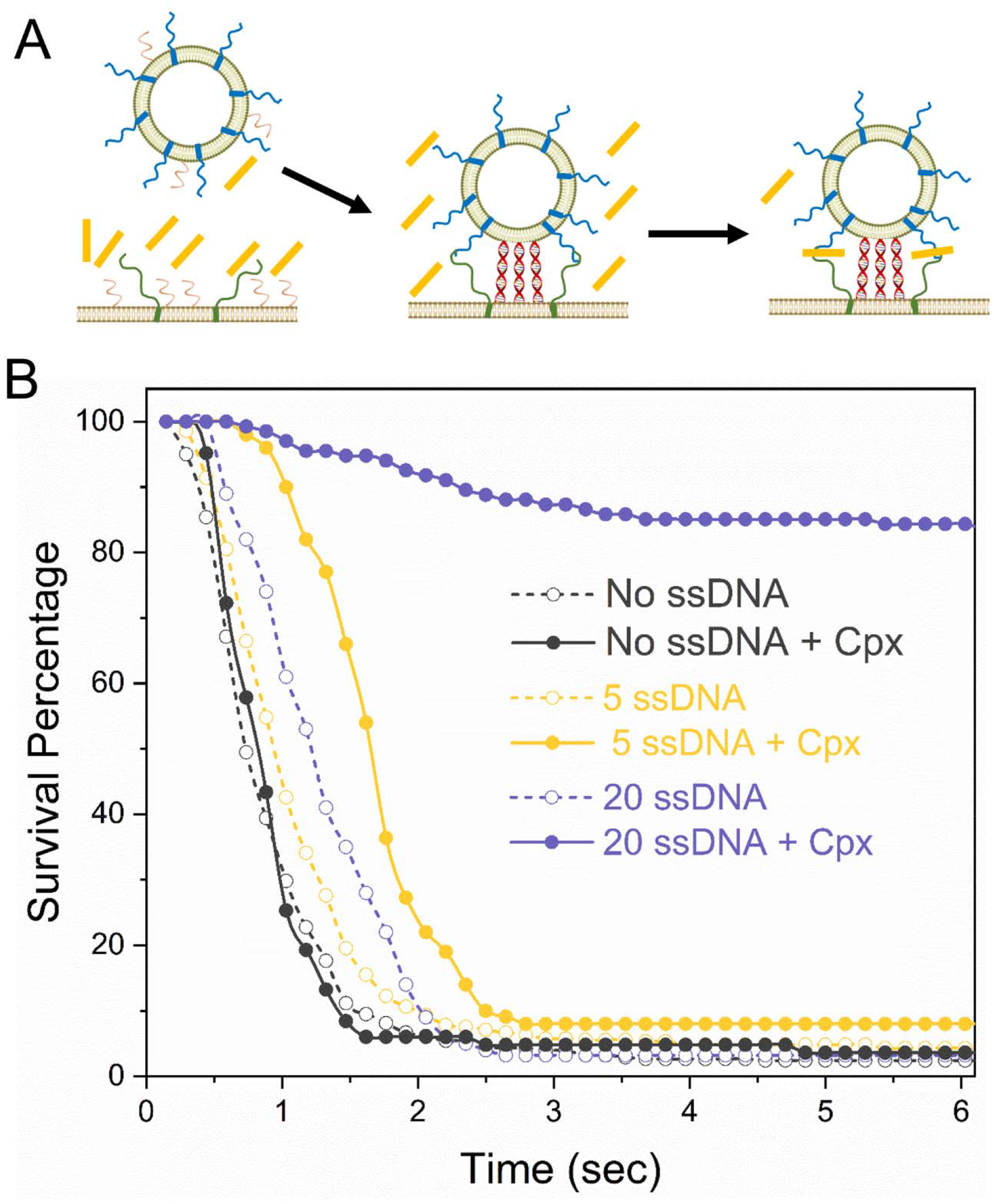
Syt1 and Cpx act sequentially to arrest SNARE-driven fusion. (A) Schematic of the programmable DNA-based mimetic used to simulate the Syt1 clamp on the SNARE-driven fusion. Annealing of the complementary ssDNA reconstituted into the SUV and the bilayer in dsDNA sterically counters the polarized SNARE assembly process and introduces a docking-to-fusion delay reminiscent of Syt1 (B) Survival analysis (Kaplan-Meier plot) curve shows that a nominal dock-to-fusion delay introduced by 20 copies of ssDNA (purple) allows Cpx to arrest spontaneous fusion of vSUVs. In contrast, no clamping was observed with 5 copies of ssDNA (yellow) which created no appreciable delay in the fusion kinetics. This suggests a sequentially mode of action for Syt1 and Cpx, wherein the kinetic delay introduced by Syt1 enables Cpx to block SNARE-driven fusion. Data was obtained from a minimum of three independent experiments, with at least 100 vesicles analyzed for each condition. A representative survival curve is shown for clarity.

We then assessed the effect of mCPX on ssDNA-regulated fusion of vSUV in the absence of Ca^2+^ (Figure 1). mCPX was able to near-completely arrest spontaneous fusion of vSUV to generate stably docked vesicles, provided that the rate of SNARE-mediated fusion was sterically delayed by ~20 copies of ssDNA (Figure 1). The majority of the vSUVs were immobile following docking to the t-SNARE-containing suspended bilayer (Figure 1), and they rarely fused over the initial observation period. In contrast, little or no inhibition was observed in control experiments with ~5 copies of ssDNA that did not introduce a detectable delay in the fusion process, as all docked vesicles proceeded to fuse spontaneously typically within 1-2 sec (Figure 1). This suggests that it is the delay in fusion per se that is necessary for the mCPX inhibitory function, and importantly that the mCPX clamp is not dependent or influenced by the ssDNA molecules. Thus, our data suggests that Syt1 and mCPX likely act sequentially to produce a synergistic clamp, with the delay introduced by Syt1 meta-stable clamp enabling CPX to bind and block the full assembly of the SNARE complex.

Next, we investigated the role of the distinct domains of mCPX in establishing the fusion block using Syt1 containing vSUV (Syt1-vSUVs). On their own, a majority (~80%) Syt1-vSUVs that docked to the t-SNARE containing bilayer surface were mobile and fused on an average 5-6 sec after docking, while a small fraction (~20%) were immobile and stably clamped (Figure 2A, B). Inclusion of 2 μM wild type mCPX (mCPX^WT^) enhanced the vesicle docking rate, with an ~5-fold increase in the total number of stably docked vesicles and >95% of Syt1-vSUVs remaining immobile post-docking (Figure 2A, B). This is consistent with our earlier findings (Ramakrishnan et al. 2020). A truncation mutant (mCPX^26-134^) lacking the unstructured N-terminal domain had very little or no effect on the vesicle docking rate or the fusion clamp, with vesicle behavior near identical to CPX^WT^ (Figure 2A, B). Deletion of the CPX_acc_ in addition to N-terminal domain (mCPX^48-134^) increased the number of docked vesicles (~4-fold) but abrogated the inhibitory function with majority of the docked vesicles proceeding to fuse spontaneously (Figure 2A, B). Targeted mutations in CPX_cen_ (R48A Y52A K69A Y70A; mCPX^4A^) that disrupt its interaction with the SNAREpins greatly reduced the stimulatory effect on vesicle docking and completely abolished the fusion clamp (Figure 2A, B). In fact, both the CPX_acc_ deletion (mCPX^48-134^) and CPX_cen_ modifications (mCPX^4A^) resulted in complete loss of mCPX inhibitory function and could not be rescued even at highest concentration (20 μM) tested (Figure 2C, Figure 2 Supplement 1). Deletion of the c-terminal domain (mCPX^26-83^) lowered the clamping efficiency (Figure 2A, B) with ~ 50% vesicles clamped under the standard experimental conditions (2 μM mCPX^26-83^). However, the inhibitory function was rescued simply by raising the concentration, and was completely restored at 20 μM mCPX^26-83^ (Figure 2C, Figure 2 Supplement 1).

**Figure 2.**
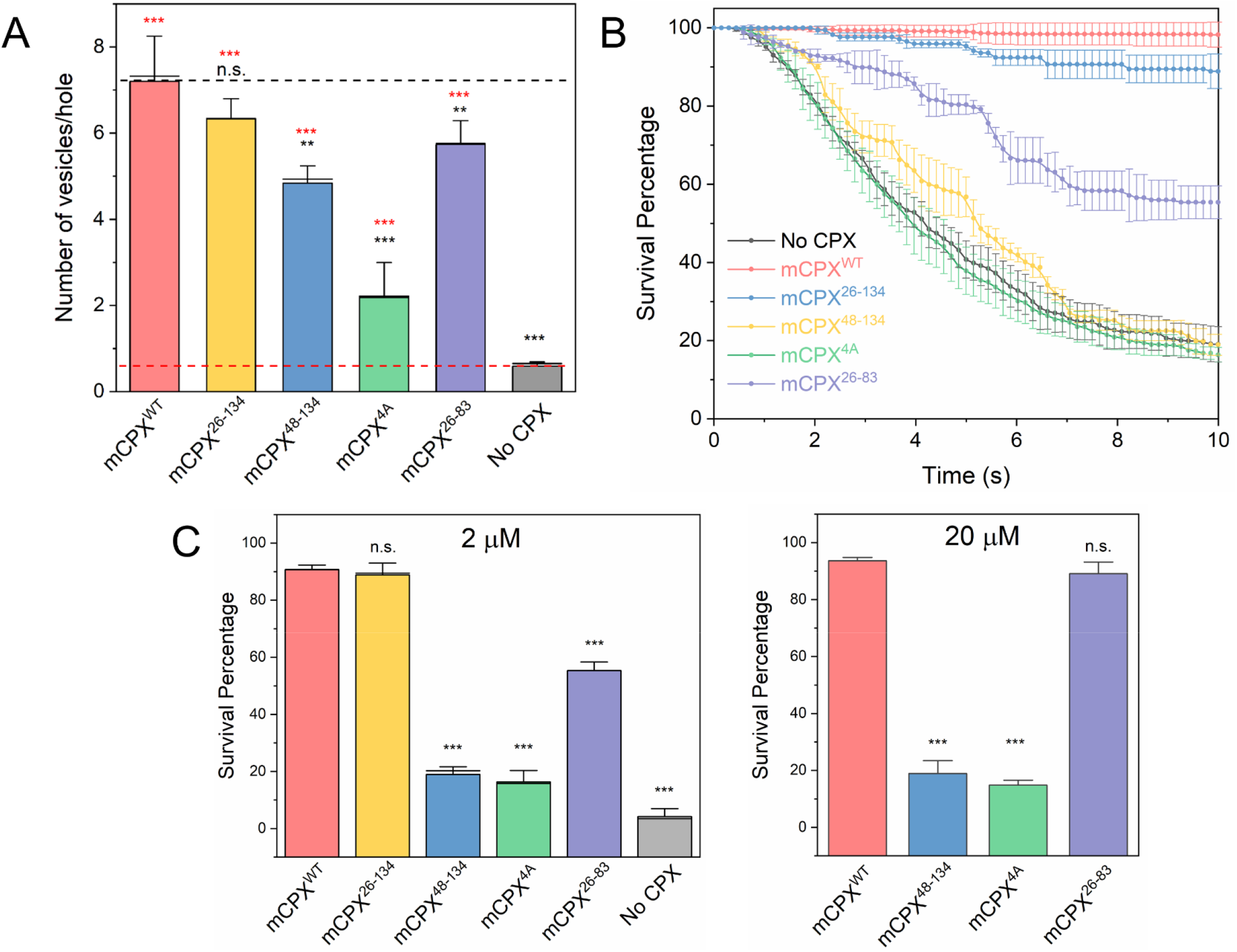
Molecular determinants of Complexin clamping function. The effect of mCPX mutants on docking and clamping of spontaneous fusion was assessed using a single-vesicle analysis with a pore-spanning bilayer setup. (A) Inclusion of mCPX increases the number of docked Syt1-vSUVs and this stimulatory effect is greatly reduced when the interaction of the CPX^cen^ to the SNAREpins is disrupted targeted mutations (mCPX^4A^). In contrast, deletion of the N-terminal domain (CPX^26-134^) or accessory helix (CPX^48-134^) or the c-terminal portion (CPX^26-83^) exhibit limited effect of the vesicle docking. In all cases, a mutant form of VAMP2 (VAMP2^4X^) which eliminated fusion was used to unambiguously estimate the number of docked vesicles after the 10 min interaction phase. (B) The time between docking and fusion was measured for each docked vesicle and the results for the whole population are presented as a survival curve (Kaplan-Meier plots). Syt1-vSUVs (black curve) are diffusively mobile upon docking and fuse spontaneous with a half-time of ~5 sec. Addition of soluble mCPX (red curve) fully arrest fusion to produce stably docked SUVs that attach and remain in place during the entire period of observation. CPX mutants with impaired SNARE interaction (mCPX^4A^, green curve) or lacking the accessory helical domain (mCPX^48-134^, yellow curve) fail to clamp fusion whilst the removal of c-terminal portion (mCPX^26-83^, purple curve) produces a partial clamping phenotype. The N-terminal domain is not involved in establishing the fusion clamp (C) End-point analysis at 10 sec post-docking shows that the both the accessory helix deletion (mCPX^48-134^) and CPXcen modifications (mCPX^4A^) result in complete loss of inhibitory function and cannot be rescued even at 20 μM concentration. In contrast, the clamping function of the c-terminal deletion mutant (mCPX^26-83^) is fully restored at high CPX concentration. The average values and standard deviations from three independent experiments (with ~300 vesicles in total) are shown. **p<0.01; *** p<0.001 using the Student’s t-test.

Altogether, we conclude that the CPX_cen_-SNAREpin interaction promotes vesicle docking, and this interaction along with CPX_acc_ are critical for mCPX mediated clamping under these physiologically-relevant experimental conditions. The c-terminal domain plays an auxiliary role and contributes to the mCPX inhibitory function likely by concentrating it on vesicle surfaces due to its curvature-binding region. Supporting this, a CPX mutant (CPX^L117W^) that enhances the curved membrane association of the c-terminal domain (Seiler et al. 2009) increased the clamping efficiency as compared to CPX^WT^ (Figure 2 Supplement 2).

Biophysical and structural studies have demonstrated that binding of the CPX_cen_ to the SNAREpins positions the CPX_acc_ to effectively block complete SNARE assembly (Kummel et al. 2011; Giraudo et al. 2008; Krishnakumar et al. 2015). While the precise mode of action is under debate, there is evidence that this involves specific interactions of CPX_acc_ with the c-terminal region of the SNAREpins (Kummel et al. 2011; Malsam et al. 2020). Critical information about these inter-molecular interactions was provided by the X-ray structure of mCPX bound to a mimetic of a pre-fusion half-zippered SNAREpins (Kummel et al. 2011). It revealed that the CPX_cen_ is anchored to one SNARE complex, while its CPX_acc_ extends away and binds to the t-SNARE in a second SNARE complex in a site normally occupied by the C-terminus of the VAMP2 helix (Kummel et al. 2011; Krishnakumar et al. 2015). This *trans* insertion model suggest a straightforward mechanism by which CPX_acc_ can block the complete assembly of the SNARE complex (Kummel et al. 2011; Krishnakumar et al. 2015).

To ascertain if the hydrophobic CPX_acc_-t-SNARE binding interfaces observed in the crystal structure are involved in clamping in our in vitro system, we tested known CPX mutants designed to either enhance (D27L E34F R37A, ‘super-clamp’ mutant mCPX^SC^) or weaken (A30E A31E L41 A44E, ‘non-clamp’ mutant 1 mCPX^NC1^) this interaction (Giraudo et al. 2009; Kummel et al. 2011). Survival analysis of Syt1-vSUVs showed that the binding interface mutants indeed alter the inhibitory activity of CPX as predicted (Figure 3A, Figure 3 Supplement 1). The mCPX^NC1^ abrogated the fusion clamp and was inactive even at higher (20 μM) concentration (Figure 3A, Figure 3 Supplement 1). In contrast, mCPX^SC^ increased the clamping efficiency and produced stably docked vesicles at lower concentrations (IC_50_ ~ 0.5 μM) compared to the mCPXWT (IC_50_ ~ 1 μM) (Figure 3B, Figure 3 Supplement 1). These findings strongly support the notion that the CPX_acc_-t-SNARE interactions observed in the pre-fusion mCPX-SNAREpin crystal is relevant for the CPX clamping function and is physiologically relevant.

**Figure 3.**
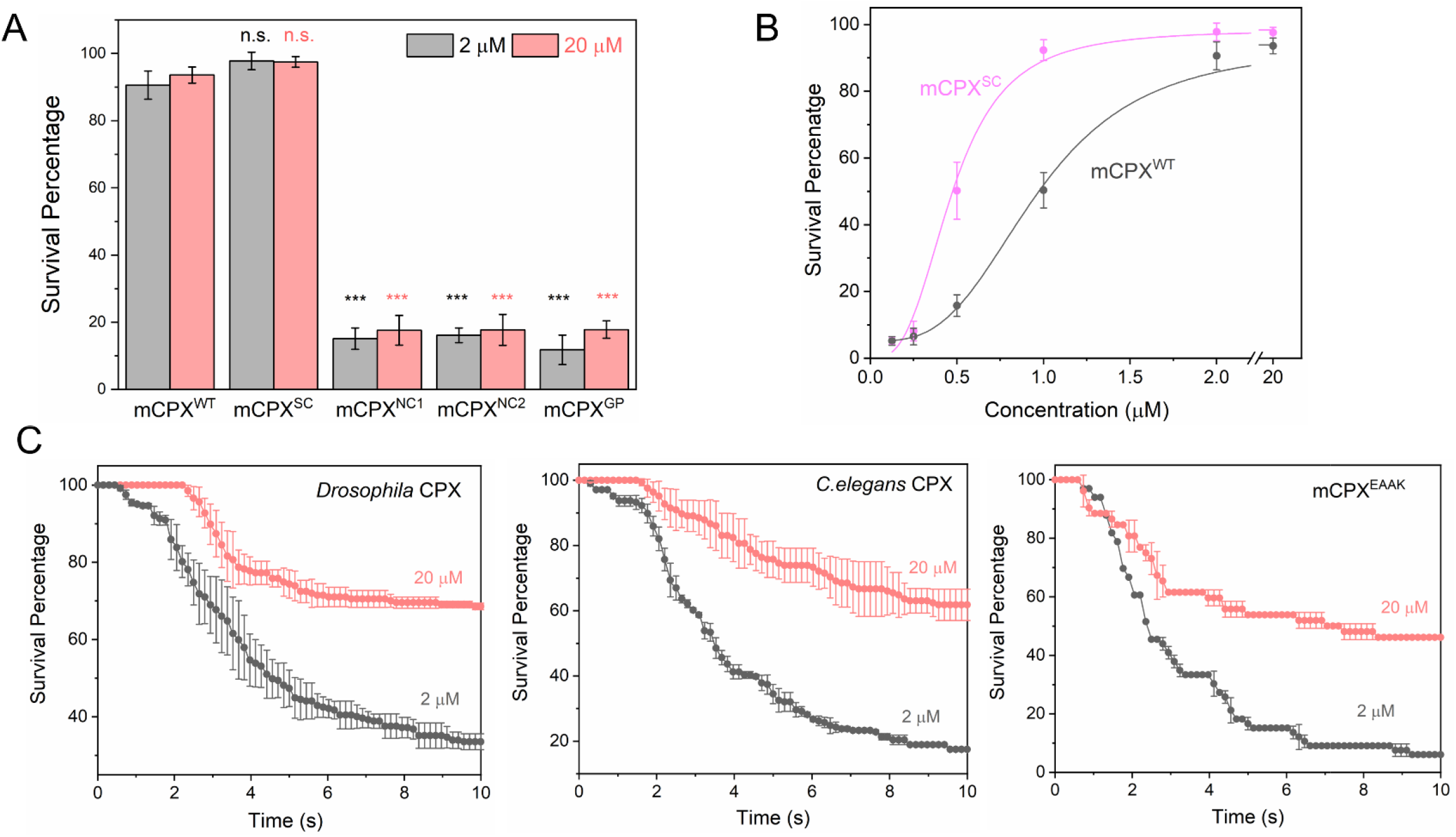
Specific interaction of mCPX accessory helix with SNAREs enhances its clamping function. (A) End-point survival analysis (measured at 10 sec post docking) using Syt1-vSUVs demonstrates that disrupting the binding of the CPX^acc^ to either the t-SNAREs (CPX^NC1^) or the VAMP2 (CPX^NC2^) abrogates the clamping function, and that a helix breaking mutation (CPX^GP^) introduced between CPX_cen_ and CPX_acc_ also abrogates the fusion clamp. (B) In contrast, mutations designed to enhance the binding of CPX^acc^ to t-SNAREs (CPX^SC^) increase the potency of the CPX clamp. Thi indicates efficient clamping by CPX requires a continuous rigid helix along with specific interaction of the CPX_acc_ with the assembling SNARE complex. (C) Supporting this notion, survival analysis (Kaplan-Meier plots) shows that both Drosophila and C. elegans CPXs, which have very low sequence identity with the mCPX accessory domain, and a CPX mutant with a randomized accessory helical sequence (CPXEAAK) have poor clamping efficiency under standard (2 μM) experimental conditions and only partial clamping at higher (20 μM) concentration. The average values and standard deviations from three to four independent experiments (with ~250 vesicles in total) are shown. *** indicates p<0.001 using the Student’s t-test.

Another key feature of the pre-fusion crystal structure is that the mCPX helix (CPX_cen_ + CPX_acc_) forms a rigid bridge between two SNARE complexes (Kummel et al. 2011; Krishnakumar et al. 2015). To test whether the rigidity of mCPX is important for clamping, we used a mCPX mutant (mCPX^GP^) having a helix-breaking linker (GPGP) inserted between CPX_cen_ and CPX_acc_. We found that disrupting the continuous helix indeed reduced the clamping efficiency (Figure 3A, Figure 3 Supplement 1) indicating that the continuity and rigidity of the CPX helix is mechanistically important for its inhibitory function.

Recently, site-specific photo-crosslinking studies in a reconstituted fusion assay revealed that CPX_acc_ (of closely related mammalian isoform CPXII) binds to the c-terminal portions of SNAP25 and VAMP2 and both interactions are important for the mCPX inhibitory function (Malsam et al. 2020). The binding interface for SNAP25 was nearly identical to CPX_acc_-t-SNARE interface observed in the crystal structure while the opposite side of the CPX_acc_ was found to interact with VAMP2 (Malsam et al. 2020). Note that this portion of VAMP2 was missing in the pre-fusion SNAREpin mimetic used for in the crystal structural analysis (Kummel et al. 2011). To understand if the aforementioned CPX_acc_-VAMP2 interaction is also part of the clamping mechanism in our cell-free system, we used a mCPX mutant (K33E R37E A40K A44E; non-clamp mutant 2, mCPX^NC2^) that reverses the charge on key binding residues and is thus expected to disrupt this interaction (Malsam et al. 2020). mCPX^NC2^ also failed to clamp spontaneous fusion of Syt1-vSUVs in our in vitro assay (Figure 3A, Figure 3 Supplement 1) and was phenotypically analogous to the t-SNARE non-binding mutant (mCPX^NC1^). This indicated the CPX_acc_ interacts with both t- and v-SNAREs to block full-zippering. As expected, because their central helix is unaltered, the majority of CPX_acc_ mutants tested retained the ability to promote vesicle docking process albeit lower than mCPX^WT^ (Figure 3 Supplement 2).

CPX_cen_ is broadly conserved (~75% amino acid sequence identity) across diverse species, whereas CPX_acc_ is highly divergent (~25% sequence identity) (Martin et al. 2011). Nonetheless, cross-species rescue experiments have been largely successful, and in fact, CPX_acc_ could be exchanged without impairing function in mammalian, fly and nematode synapses (Xue et al. 2009; Cho et al. 2014; Radoff et al. 2014). This raises the question whether the distinct CPX_acc_-SNARE interactions that are vital for mCPX inhibitory functionality in our *in vitro* assays are physiologically relevant. To address this, we examined the clamping ability of the *C. elegans* (ceCPX) and *Drosophila* (dmCPX) orthologs of mCPX in our *in vitro* reconstituted assay. Under standard experimental conditions (2 μM CPX), both ceCPX and dmCPX were able to promote vesicle docking (Figure 3 Supplement 2) but were considerably less efficient (~15% and ~30% respectively) in preventing spontaneous fusion of Syt1-vSUV (Figure 3C) as compared near-complete (>95%) fusion clamp observed with mCPX (Figure 2B, C). However, simply increasing the concentrations improved the clamping efficacy of both dmCPX and ceCPX, with ~60-70% of docked vesicles stably-clamped at 20 μM concentration (Figure 3C) and remained Ca^2+^-sensitive (Figure 3 Supplement 3).

This suggests that specific molecular interactions of CPX_acc_ with SNAREs likely increase the potency of the mCPX inhibitory function and that this effect may be occluded at high concentrations of CPX. To verify this, we examined the effect of the mCPX mutant wherein the endogenous CPX_acc_ domain is replaced with an artificial alpha helix based on a Glu-Ala-Ala-Lys (EAAK) motif repeated seven times (Radoff et al. 2014). Noteworthy, this construct (mCPX^EAAK^), which contains similar charge/hydrophobic residues but in random order with an overall ~30% sequence identity to CPX_acc_, was able to fully-restore CPX inhibitory functionality in *C. elegans* neuromuscular synapses (Radoff et al. 2014). In our *in vitro* assay, CPX^EAAK^ enhanced initial docking (Figure 3 Supplement 2) but failed to clamp spontaneous fusion (~10% efficiency) under standard experimental conditions (2 μM CPX) and was moderately effective (~50% efficiency) at higher (20 μM CPX) concentration (Figure 3C, Figure 3 Supplement 3). This supports the notion the specific CPX_acc_-SNARE interaction is functionally-relevant and likely enhances CPX inhibitory function.

Finally, we evaluated the probability and rate of Ca^2+^-triggered fusion from the clamped state in the presence and absence of mCPX. We used Syt1-vSUV loaded with Sulforhodamine B (fluorescent content marker) to track full-fusion events and lipid-conjugated Ca^2+^ indicator (Calcium green C24) attached to the planar bilayer to estimate the time of arrival of Ca^2+^ at/near the docked vesicles (Figure 4). Consistent with our previous study, the influx of free Ca^2+^ (1 mM) triggered simultaneous fusion of >90% of the Syt1/CPX-clamped vesicles (Figure 4). These vesicles fused rapidly and synchronously, with a characteristic time-constant (τ) of ~10 msec following the arrival of Ca^2+^ locally (Figure 4). Considering that the majority of Ca^2+^-triggered fusion occurs within a single frame (13 msec), we suspect that the true Ca^2+^-driven fusion rate is likely <10 msec (Figure 4).

**Figure 4.**
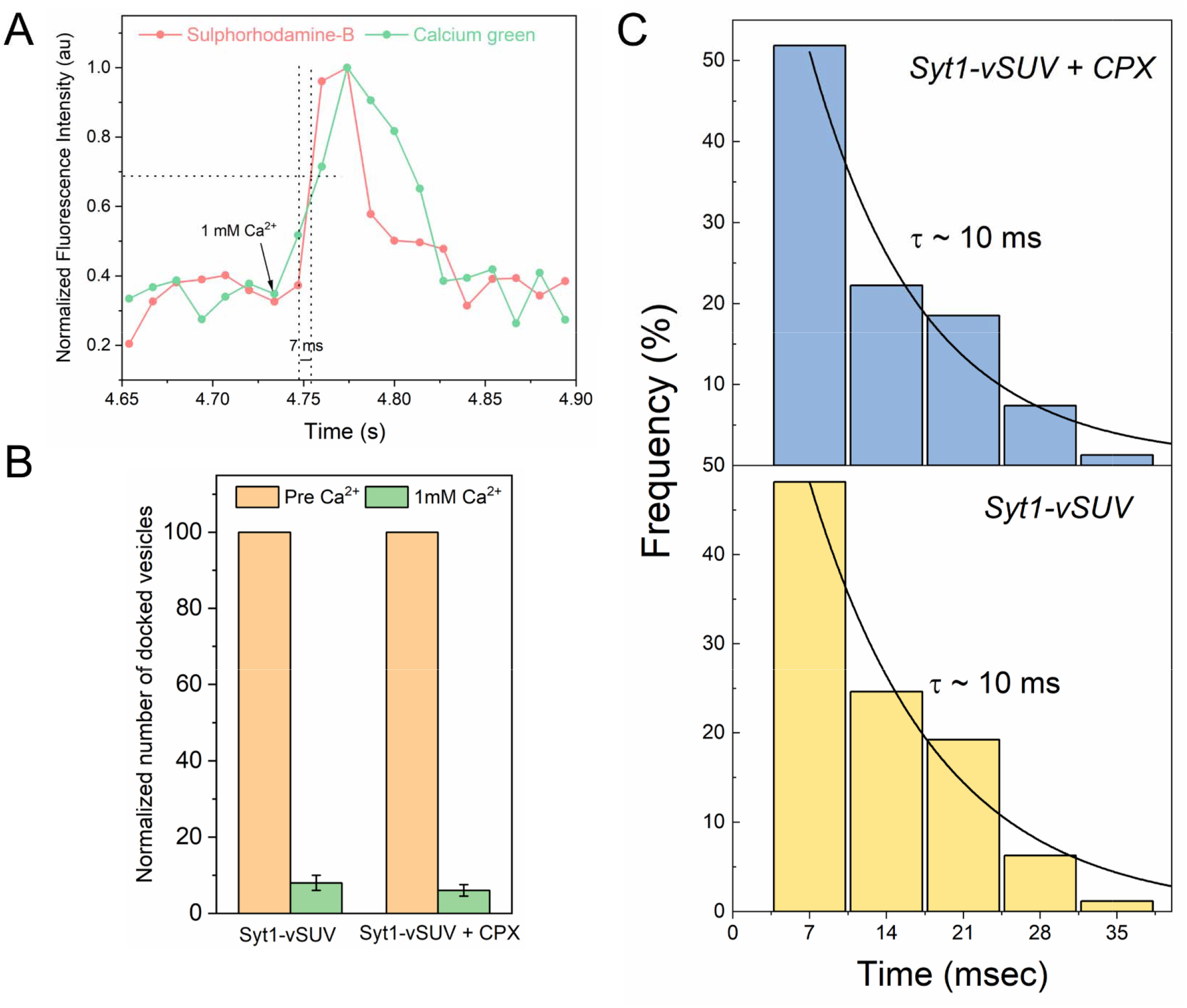
Complexin does not contribute to the probability or speed of Ca^2+^-triggered vesicular release. (A) The effect of CPX on Ca^2+^-triggered fusion was assessed using a content-release assay with Sulforhodamine B loaded vesicles. Sulforhodamine B is largely self-quenched when encapsulated inside an SUV. Fusion of the vesicle results in dilution of the probe, which is accompanied by increasing fluorescence. The Ca^2+^-sensor dye, Calcium Green, introduced in the suspended bilayer (via a lipophilic 24-carbon alkyl chain) was used to monitor the arrival of Ca^2+^ at/near the docked vesicles. A representative fluorescence trace before and after the addition of 1 mM Ca^2+^ shows that the rise in Sulforhodamine B (red curve) fluorescence intensity occurs within a single frame (13 ms) of Ca^2+^ binding to local Calcium green (green curve) (B) End-point analysis at 1 min post Ca^2+^-addition shows that >90% of all clamped vesicles fuse following Ca + addition both in the presence or absence of CPX. (C) Kinetic analysis shows that the clamped vesicles with or without CPX fuse rapidly following Ca^2+^-addition with near identical time constant of ~10 msec. This represents the temporal resolution limit of our recordings (13 ms frame rate) and the true Ca^2+^-triggered fusion rate may well be below 10 msec. In all cases, the effect of CPX was assessed under physiologically-relevant SUV conditions (~74 copies of VAMP2 and ~25 copies of Syt1), while vesicles containing low copy VAMP2 (~13 copies) with normal number (~25 copies) of Syt1 molecules was used in the absence of CPX. The average values and standard deviations from three independent experiments (with ~200 vesicles in total) are shown.

In absence of mCPX, we observed a relatively small number of docked vesicles prior to Ca^2+^-influx and this precluded any meaningful quantitative analysis. Hence, to obtain stably docked vesicles without mCPX, we used low VAMP2 conditions i.e. SUVs containing ~13 copies of VAMP2 and ~25 copies of Syt1. We have previously demonstrated that under these conditions, Syt1 alone is sufficient to produce stably-clamped vesicles (Ramakrishnan et al. 2019) and that is what we observed with >95% of docked vesicles immobile post-docking. Addition of Ca^2+^ (1 mM) triggered rapid and synchronous fusion of >90% of all docked vesicles, with τ ~ 10 msec. Indeed, these vesicles indistinguishable in their behavior compared to the physio-mimetic SUVs in the presence of mCPX (Figure 4) indicating the mCPX is not critical for Ca^2+^-activation mechanism and Syt1 is sufficient to achieve Ca^2+^-synchronized rapid vesicular fusion.

## DISCUSSION

There is a long-standing debate over the role of CPX in establishing a fusion clamp and perhaps the best evidence in support has come from biochemical analyses (Giraudo et al. 2006; Giraudo et al. 2008; Kummel et al. 2011; Lai et al. 2014) and physiological studies in invertebrate synapses (Huntwork and Littleton 2007; Cho et al. 2014; Martin et al. 2011; Hobson et al. 2011). In the case of mammalian synapses, a role for CPX in blocking spontaneous release events remains controversial because KD/KO manipulations yield seemingly contradictory results and show neuron-specific differences (Xue et al. 2008; Lopez-Murcia et al. 2019; Maximov et al. 2009; Yang et al. 2013). Here, using a fully defined albeit simplified cell-free system we provide compelling evidence that mCPX is an integral part of the overall clamping mechanism and delineate the molecular determinants of mCPX inhibitory function. The distinct effects of different CPX truncation and targeted mutations match with data obtained from other reductionist or even physiological systems (Giraudo et al. 2006; Giraudo et al. 2008; Kummel et al. 2011; Cho et al. 2014; Lai et al. 2014; Gong et al. 2016) forcefully arguing for the physiological relevance of results obtained from our in vitro reconstituted assay.

Taken together, our experiments indicate that mCPX inhibitory function entails distinct and specific interactions of the CPX_cen_ and CPX_acc_ domains with assembling SNAREpins, and that the c-terminal domain augments clamping function by increasing the local concentration and/or by proper orientation of CPX via interactions with the vesicle membrane (Figure 2). Our results indicate that CPX_cen_ binds in the groove between assembling Syntaxin and VAMP2 helices at the early stages of vesicle docking to stabilize the partially-zippered SNAREpins, consequently promote vesicle docking. This in turn positions CPX_acc_ to block further zippering of SNARE complex both by directly capturing the VAMP2 c-terminus and by simultaneously occupying its binding pocket on the t-SNARE. The precise configuration of the clamped state under the docked vesicles remains to the determined, but we find that a continuous, rigid CPX helix is essential for a stable fusion clamp. The only plausible way for both CPX_cen_ and CPX_acc_ to interact with the SNAREpins is if CPX interacts with two different pre-fusion SNAREpins as observed in the pre-fusion CPX-SNARE X-ray crystal structure (Kummel et al. 2011). Thus, our data strongly support the *trans* insertion clamping model specifically for the peripheral SNAREs that uniquely require CPX for clamping in the reconstituted system (Ramakrishnan et al. 2020), wherein CPX clamps SNARE-mediated fusion by blocking complete zippering of the VAMP2 on a neighboring SNARE complex.

Noteworthy, we observe that the specific interactions of the CPX_acc_ with the synaptic SNARE proteins increase the potency of the clamp, and in accordance mCPX is ~2-3 fold more efficient in establishing the fusion clamp as compared to dmCPX or ceCPX under the same experimental conditions (Figure 3). However, the divergence in clamping ability among the mammalian, fly and nematode CPXs is diminished at higher concentrations of CPX. This might explain the puzzling fact that in physiological analyses, when CPX is over-expressed, cross-species rescue experiments are largely successful yet CPX_acc_-SNARE disrupting mutants’ exhibit limited effect on the CPX clamping ability (Yang et al. 2010; Cho et al. 2014; Radoff et al. 2014). Considering that the CPX_acc_ is highly divergent across different species, it is conceivable that CPX_acc_ has distinctively evolved to optimally bind and clamp the species-specific SNARE partners. Additional biochemical/structural studies are needed to address this question.

Overall, our data strongly argues that mCPX has an intrinsic ability to inhibit SNARE-dependent fusion and under minimal conditions is required, along with Syt1, to generate and maintain a pool of release-ready vesicles. Indeed, functionality of mCPX observed in our *in vitro* system perfectly matches with physiological studies in model invertebrate systems (Martin et al. 2011; Hobson et al. 2011; Huntwork and Littleton 2007; Cho et al. 2014). However, recent physiological studies in mammalian synapses reported that acute CPX loss reduces SV fusion probability but does not unclamp spontaneous fusion. Hence, they conclude that CPX is dispensable for ‘fusion clamping’ in mammalian neurons (Lopez-Murcia et al. 2019). It is worth noting that under these conditions CPX removal abates both spontaneous and evoked neurotransmitter release without changing the number of docked vesicles (Lopez-Murcia et al. 2019). This suggests that acute CPX loss likely affects the late-stage vesicle priming process, and it is possible this ‘loss-of-fusion’ phenotype occludes CPX role in regulating spontaneous fusion events. It is also feasible mCPX plays a more specialized role in mammalian synapses and is primarily involved in stabilizing newly primed synaptic vesicles and prevents their premature fusion (Dhara et al. 2014; Chang et al. 2015). Thus, mCPX functions as a fusion clamp in an activity-dependent manner and is critical to blocking spontaneous/tonic and asynchronous vesicular release (Dhara et al. 2014; Chang et al. 2015; Yang et al. 2010), and thus, promoting synchronous SV exocytosis.

mCPX on its own is ineffective in clamping SNARE-driven vesicle fusion, as the c-terminal portion of VAMP2 assembles into the SNARE complex far faster than free CPX can bind to prevent its zippering (Gao et al. 2012). As such, a delay in SNARE zippering is required for the CPX to bind and thereby block fusion. The fact that sufficient delay can be artificially provided by ~20 copies of DNA duplexes (Figure 1) suggest that under physiological conditions, Syt1 (and perhaps other proteins on the SV) might hinder the SNARE assembly by a simple steric mechanism, enabling mCPX to function as a fusion clamp. This is supported by the observation that the Syt1 clamp or the formation of the central SNAREpins are not strictly required for mCPX clamping function (Ramakrishnan et al. 2020).

Ca^2+^-activation studies (Figure 4) show that mCPX has no observable effect of the probability or speed of vesicle fusion from the clamped state. Indeed, the probability and speed of vesicle fusion from the clamped state were identical whether the vesicles were clamped by both Syt1/Cpx (i.e. containing both central and peripheral SNAREpins) *or* only by Syt1-associated central SNAREpins (Figure 4). Evidently, the activation of the Syt1-central SNAREpins are sufficient to achieve Ca^2+^-regulated release, at least with our current time resolution of ~ 13 msec and high Ca^2+^ levels

Besides clamping the Syt1-independent peripheral SNAREpins, it could also bind the Syt1-associated central SNAREpin (Zhou et al. 2017). In fact, recent crystal structure shows that mCPX binding creates a new binding interface for second Syt1 to bind the same SNARE complex (Zhou et al. 2017). Our earlier work indicated that this ‘tripartite’ interface is not necessary to produce stably docked vesicles, but likely required for efficient Ca^2+^-triggered fusion from the clamped state (Ramakrishnan et al. 2020). So it is conceivable that mCPX indirectly regulates Ca^2+^ synchronized vesicular fusion via the ‘tripartite’ interface. Furthermore, as the tripartite binding motif is largely conserved among different Syt isoforms, it is possible that mCPX binding could enable synergistically regulation of vesicular release by different calcium sensors (Volynski and Krishnakumar 2018). As such, further studies with high temporal resolution, physiological Ca^2+^ dynamics and different calcium sensors is needed to dissect the precise role of mCPX in modulating Ca^2+^-activation of fusion from the clamped state.

## MATERIALS AND METHODS

### Proteins and Materials

The following cDNA constructs, which have been previously described (Krishnakumar et al. 2013; Ramakrishnan et al. 2019; Ramakrishnan et al. 2020), were used in this study: full-length VAMP2 (VAMP2-His^6^, residues 1-116); full-length VAMP2^4X^ (VAMP2-His^6^, residues 1-116 with L70D, A74R, A81D, L84D mutations), full-length t-SNARE complex (mouse His^6^-SNAP25B, residues 1-206 and rat Syntaxin1A, residues 1-288); Synaptotagmin (rat Synaptotagmin1-His^6^, residues 57-421); and Complexin (human His^6^-Complexin 1, residues 1-134). All CPX mutants (truncations/point-mutations) were generated in the same background. All proteins were expressed and purified as described previously (Krishnakumar et al. 2013; Ramakrishnan et al. 2019; Ramakrishnan et al. 2020). All the lipids used in this study were purchased from Avanti Polar Lipids (Alabaster, AL). ATTO647N-DOPE was purchased from ATTO-TEC, GmbH (Siegen, Germany) and Calcium Green conjugated to a lipophilic 24-carbon alkyl chain (Calcium Green C24) was custom synthesized by Marker Gene Technologies (Eugene, OR). HPLC-purified DNA sequences (5’-ATCTCAATTATCCTATTAACC-3’ and 5’-GGTTAATAGGATAATTGAGAT-3’) conjugated to cholesterol with a 15 atom triethylene glycol spacer (DNA-TEG-Chol) were synthesized at Yale Keck DNA sequencing facility.

### Liposome Preparation

VAMP2 (± Syt1) were reconstituted into small unilamellar vesicles (SUV) were using rapid detergent (1% Octylglucoside) dilution and dialysis method as described previously (Ramakrishnan et al. 2019; Ramakrishnan et al. 2020). The proteo-SUVs were further purified via float-up using discontinuous Nycodenz gradient. The lipid composition was 88 (mole) % DOPC, 10% PS and 2% ATTO647-PE for VAMP2 (± Syt1) SUVs and we used protein: lipid (input) ratio of 1:100 for VAMP2 for physiological density, 1: 500 for VAMP2 at low copy number, and 1: 250 for Syt1. Based on the densitometry analysis of Coomassie-stained SDS gels and assuming the standard reconstitution efficiency, we estimated the vesicles contain 73 ± 6 (normal physiological-density) or 13 ± 3 (low-density) and 25 ± 4 copies of outward-facing VAMP2 and Syt1 respectively (Figure 1 Supplement 1).

### Single Vesicle Fusion Assay

All the single-vesicle fusion measurements were carried out with suspended lipid bilayers as previously described (Ramakrishnan et al. 2018; Ramakrishnan et al. 2019; Ramakrishnan et al. 2020). Briefly, t-SNARE-containing giant unilamellar vesicles (80 % DOPC, 15% DOPS, 3% PIP2 and 2% NBD-PE) were prepared using the osmotic shock protocol and bursted onto Si/SiO2 chips containing 5 μm diameter holes in presence of HEPES buffer (25 mM HEPES, 140 mM KCl, 1mM DTT) supplemented with 5 mM MgCl_2_. The free-standing lipid bilayers were extensively washed with HEPES buffer containing 1 mM MgCl_2_ and the fluidity of the t-SNARE containing bilayers was verified using fluorescence recovery after photo-bleaching using the NBD fluorescence.

Vesicles (100 nM lipids) were added from the top and allowed to interact with the bilayer for 10 minutes. The ATTO647N-PE fluorescence introduced in the vesicles were used to track vesicle docking, post-docking diffusion, docking-to-fusion delays and spontaneous fusion events. The time between docking and fusion corresponded to the fusion clamp and was quantified using a ‘survival curve’ whereby delays are pooled together, and their distribution is plotted in the form of a survival function (Kaplan-Meier plots). For the end-point analysis, the number of un-fused vesicles (survival percentage) was estimated ~10 sec post-docking. After the initial 10 min, the excess vesicles were removed by buffer exchange (3x buffer wash) and 1 mM CaCl_2_ was added from the top to monitor the effect of Ca^2+^ on the docked vesicles. The number of fused (and the remaining un-fused) vesicles was estimated (end-point analysis) ~1 min after Ca^2+^-addition. CPX protein (at the indicated final concentration) were added to the experimental chamber and incubated for 5 min prior to the addition of the vesicles. All experiments were carried out at 37ºC using an inverted laser scanning confocal microscope (Leica-SP5) and the movies were acquired at a speed of 150 ms per frame, unless noted otherwise. Fate of each vesicles were analyzed using our custom written MATLAB script described previously (Ramakrishnan et al. 2018). The files can be downloaded at: https://www.mathworks.com/matlabcentral/fileexchange/66521-fusion-analyzer-fas.

### Single-Vesicle Docking Analysis

To get an accurate count of the docked vesicles, we used VAMP2 mutant protein (L70D, A74R, A81D & L84D; VAMP2^4X^) that eliminates fusion without impeding the docking process (Krishnakumar et al. 2013). For the docking analyses, 100 nM VAMP2^4X^ containing SUVs (vSUV^4X^) were introduced into the chamber and allowed to interact with the t-SNARE bilayer for 10 mins. The bilayer was then thoroughly washed with the running buffer (3x minimum) and the number of docked vesicles were counted, using Image J software.

### DNA-regulated Single Vesicle Fusion Assay

To prepare ssDNA containing vesicles, dialyzed VAMP2 or t-SNARE containing SUVs were incubated with complementary DNA-TEG-Chol for 2h at room temperature with mild-shaking. The v-SUVs were further purified using the Nycodenz gradient. We used the lipid: DNA-TEG-Chol input ratios of 1:2000, 1:1000, 1:500, and 1: 200 produce vSUVs with approximately 5, 10, 20, 50 copies of ssDNA per vesicles respectively. To identify the optimal condition for the single-vesicle fusion assays, we first tested the fusogenicity of ssDNA containing vesicles using bulk-fusion assay (Figure 1 Supplement 2). Fusion of vSUV with t-SNARE liposomes were un-affected up to 20 copies of ssDNA, but we observed some reduction in fusion levels with 50 copies of ssDNA (Figure 1 Supplement 2). Correspondingly, in the single-vesicle fusion setup, vSUV with 5, 10 & 20 copies of ssDNA docked and fused spontaneously with progressive docking-to-fusion delays, but the majority of 50 ssDNA-vSUV remained docked and un-fused (Data not shown). So, we chose to test the effect of Cpx on 20 ssDNA-vSUV, with 5 ssDNA-vSUV as the control.

### Calcium Dynamics

We used a high affinity Ca^2+^-sensor dye, Calcium Green (K_d_ of ~75 nM) conjugated to a lipophilic 24-carbon alkyl chain (Calcium Green C24) introduced in bilayer to monitor the arrival of Ca^2+^. To estimate the arrival of Ca^2+^ at or near the docked vesicle precisely, as indicated by increased in Calcium green fluorescence at 532 nm, we used resonant scanner to acquire movies at a speed of up to 13 msec per frame with 512 x 32 resolution. For each vesicle fusion kinetics, calcium arrival was monitored over area of an individual hole (5 μm diameter) to get the high signal to noise ratio and vesicle fusion was monitored with 0.5 μm ROI around the docked vesicle. In these experiments, we used Sulforhodamine-B loaded Syt1-vSUV and tracked full-fusion events using increase in fluorescence signal due to dequenching of Sulforhodamine-B.

## ACKNOWLEDGEMENTS

This work was supported by National Institute of Health (NIH) grant DK027044.

---

**Figure 1: Supplement 1.**
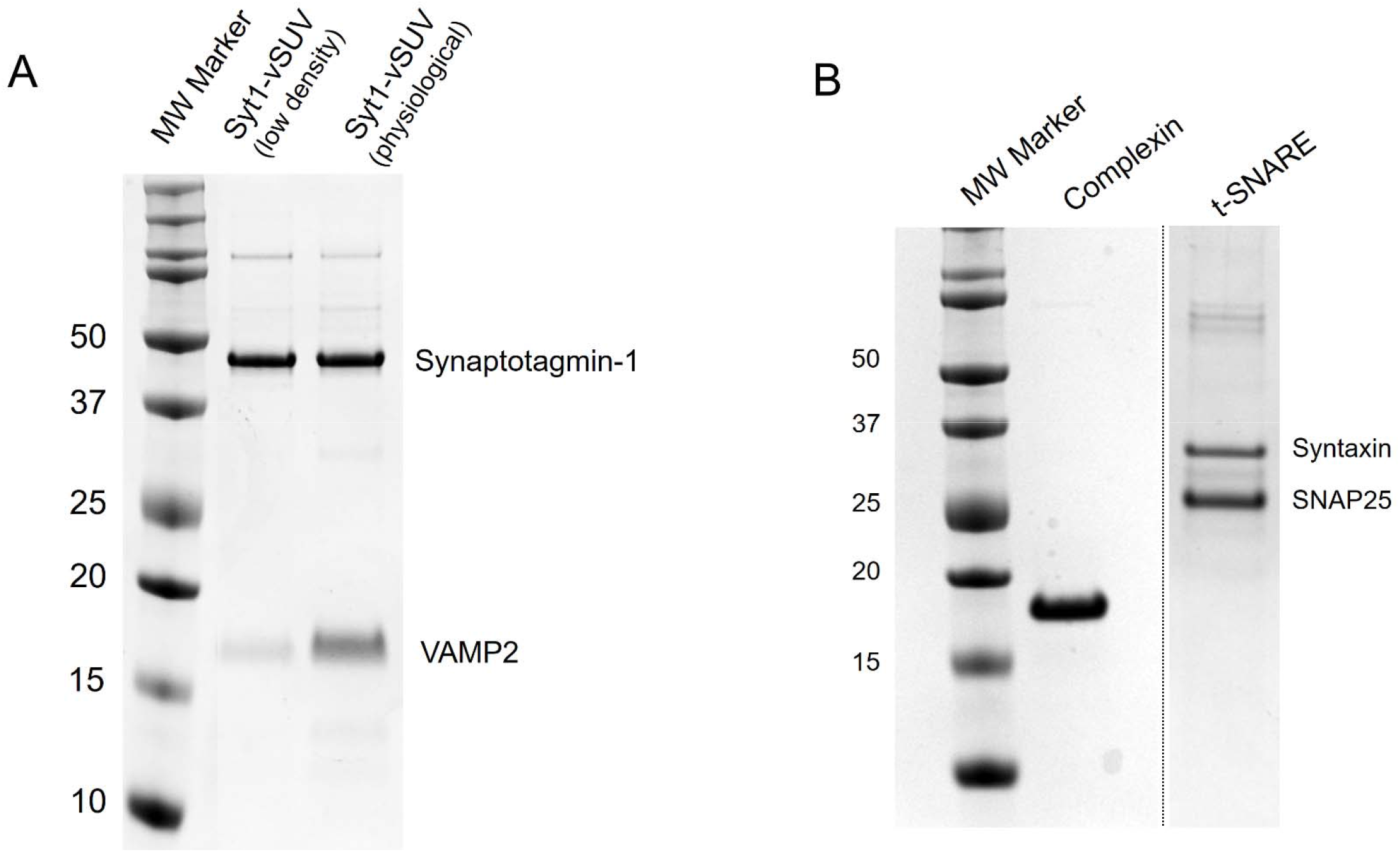
Coomassie-stained SDS-PAGE analysis of the proteins used in this study. (A) Gel image of the Syt1-VAMP2 containing SUVs used a synaptic vesicle mimic. To approximate the protein density on SVs, in most experiments we used SUVs containing 73 ± 6 and 25 ± 4 copies of outward-facing VAMP2 and Syt1. We used SUV containing 13 ± 4 and 25 ± 4 copies of VAMP2 and Syt1 under low-copy conditions. (B) In all fusion experiments, co-purified t-SNAREs containing 1:1 complex of Syntaxin1a and SNAP25 was reconstituted into the free-standing bilayer and full-length Complexin 1 (WT or mutants) were added in solution at the concentration indicated.

**Figure 1: Supplement 2.**
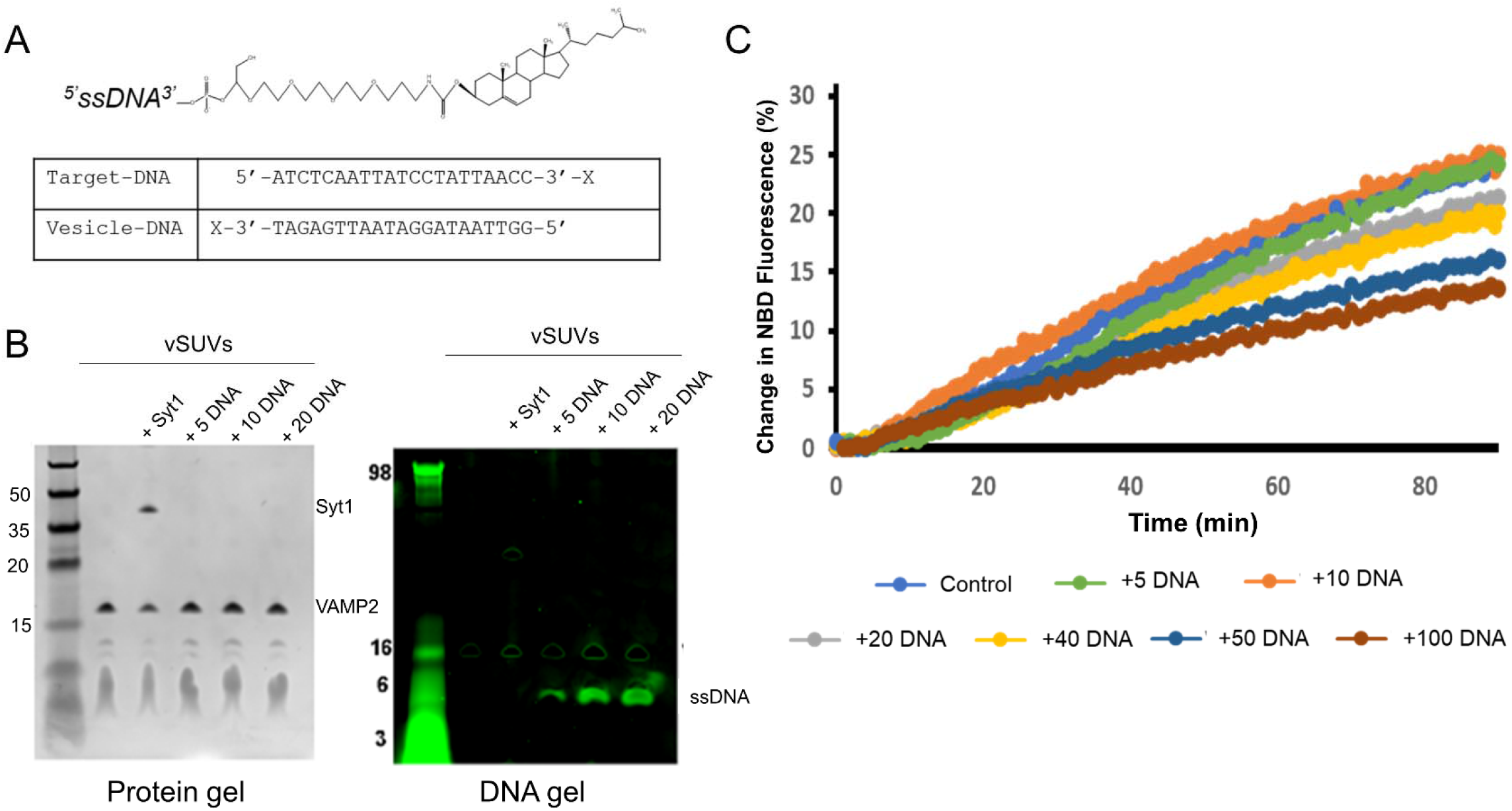
DNA-hybridization regulates SNARE-mediated membrane fusion. (A) Complementary single stranded DNA (ssDNA) sequences were conjugated to cholesterol with tri-ethylene glycol spacer were incubated with pre-formed VAMP2 or t-SNARE SUVs with mild agitation and subsequently purified using float-up with a discontinuous Nycodenz gradient (B) Representative Coomassie and Sybr-green stained gels showing the incorporation of defined number of ssDNA into v-SUVs. The number of ssDNA incorporated into the vSUVs were controlled using a defined input ssDNA: lipid ratio. (C) Bulk lipid mixing assay (Weber et al. 1998) with ssDNA incorporated v- and t-SUVs shows that SNARE-driven fusion is not appreciably affected up to relatively high ssDNA density (>50 copies of ssDNA per vesicle). Representative fusion curves measured by dequenching of NBD-fluorescence is shown for clarity.

**Figure 2: Supplement 1.**
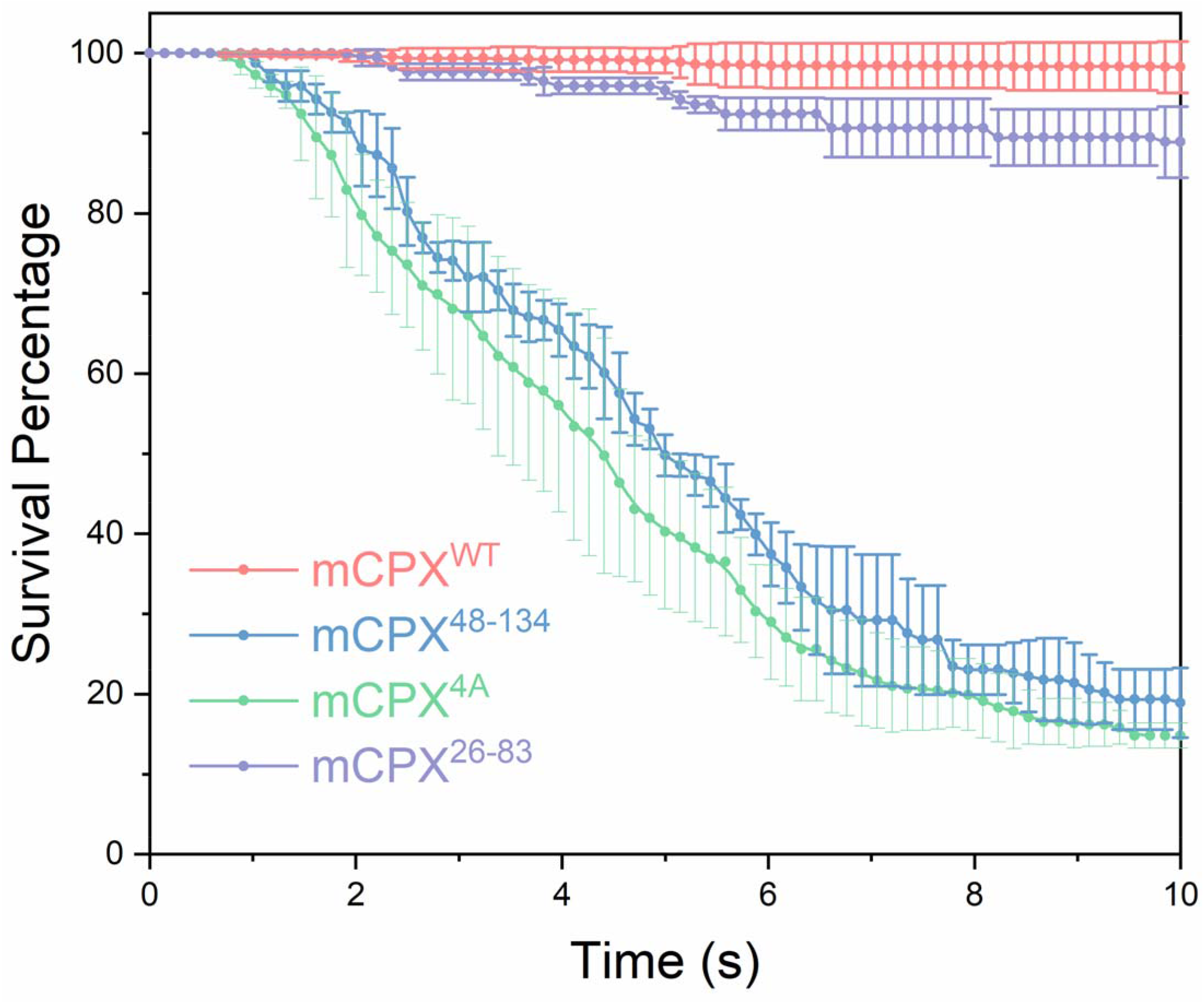
Survival analysis (Kaplan-Meier plots) of Syt1-vSUVs shows that the loss of clamping phenotypes observed with CPX mutant with impaired SNARE interaction (mCPX^4A^, green curve) or lacking the accessory helical domain (mCPX^48-134^, blue curve) is not rescued at high (20 μM) CPX concentrations. In contrast, increasing the CPX concentration fully-restores the inhibitory function of the c-terminal deletion mutant (mCPX^26-83^, purple curve). The average values and standard deviations from three independent experiments (with ~300 vesicles in total) are shown.

**Figure 2: Supplement 2.**
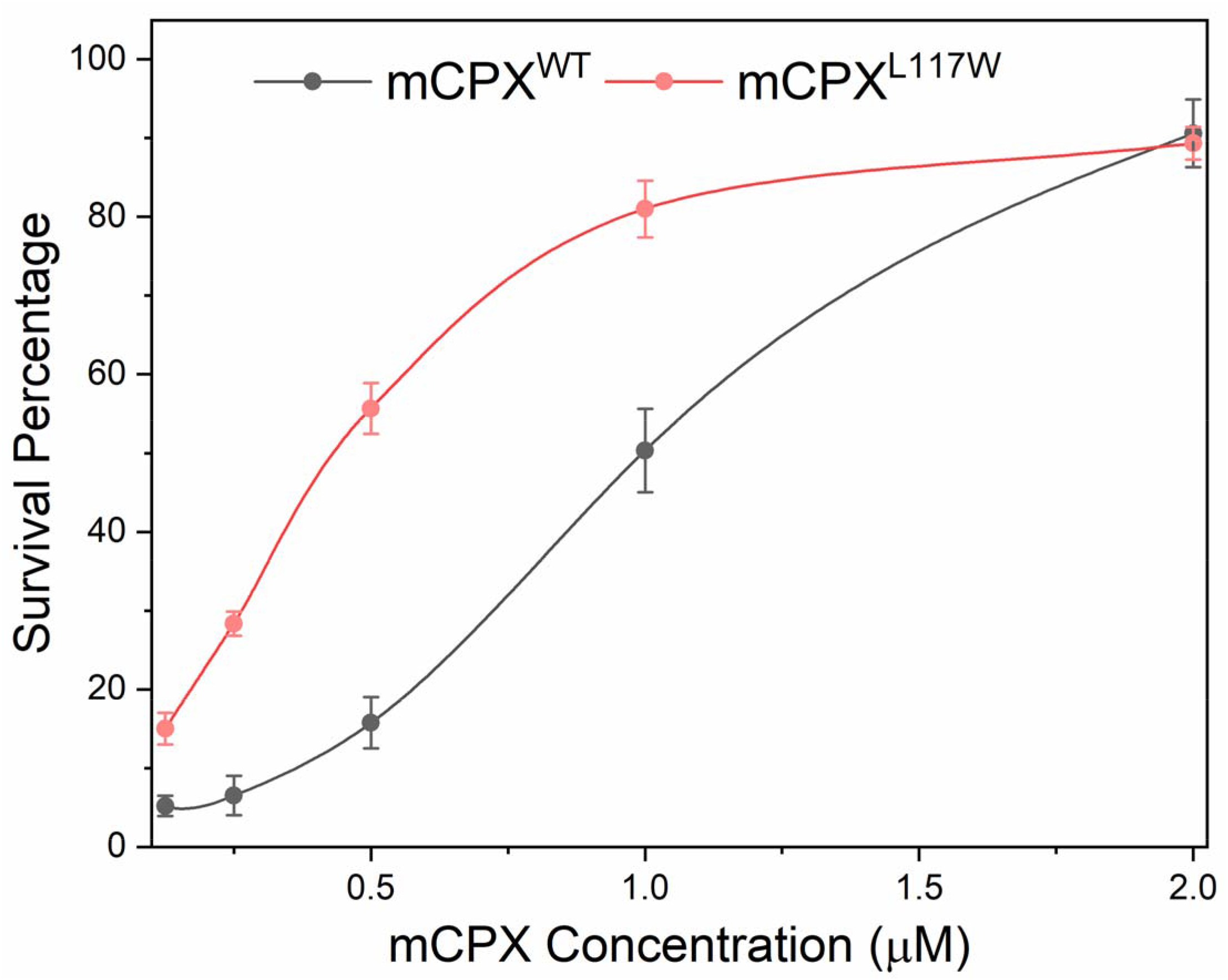
**B** Dose-dependency analysis using Syt1-vSUVs shows that CPX mutant with a hydrophobic mutation (mCPX^L117W^, red curve) designed to improve its membrane association is more efficient in clamping fusion as compared to the CPX^WT^ (black curve). This implies that the c-terminal domain contributes to clamping function by increasing the local CPX concentration via membrane interaction. The average values and standard deviations from three to four independent experiments (with ~250 vesicles in total) are shown.

**Figure 3: Supplement 1.**
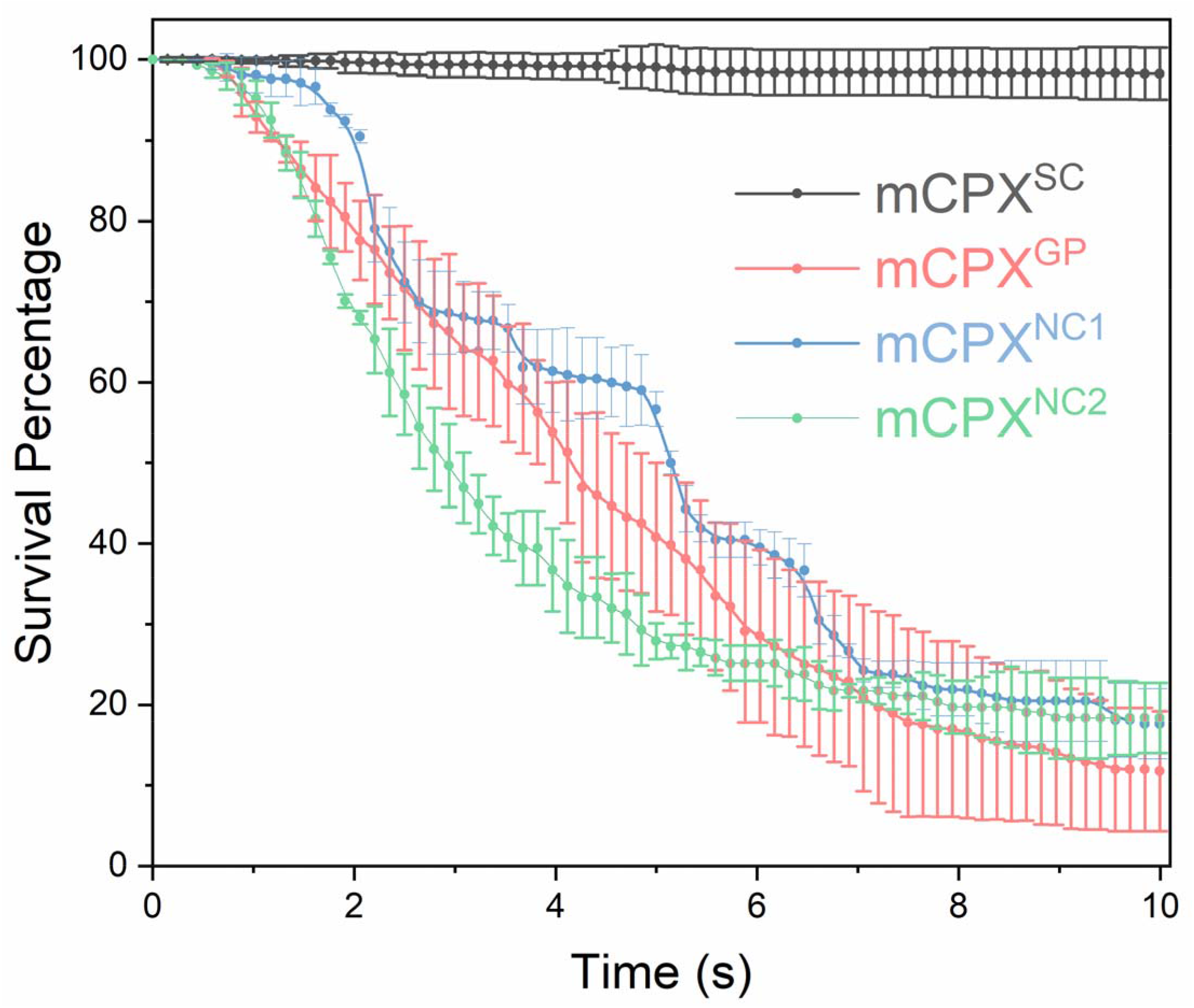
Survival analysis (Kaplan-Meier plots) using Syt1-vSUVs and 2 μM CPX shows targeted mutations that disrupt the interaction of CPX_acc_ with the t-SNARE (mCPX^NC1^, blue curve) or VAMP2 (mCPX^NC2^, green curve) abrogate the clamping function. Similarly, GPGP mutations introduced between the CPX_cen_ and CPX_acc_ (mCPX^GP^, red curve) that breaks the continuity of the helical domain disrupts the clamping function. Overall, this argues that a rigid α-helical structure along with distinct interaction of the CPX_acc_ with the assembling SNAREpins are required for its clamping function. Supporting this, CPX_acc_ that strengths the t-SNARE binding stabilizes the fusion clamp and as such, enhances the clamping efficiency (Figure 2B). The average values and standard deviations from three independent experiments (with ~300 vesicles in total) are shown.

**Figure 3: Supplement 2.**
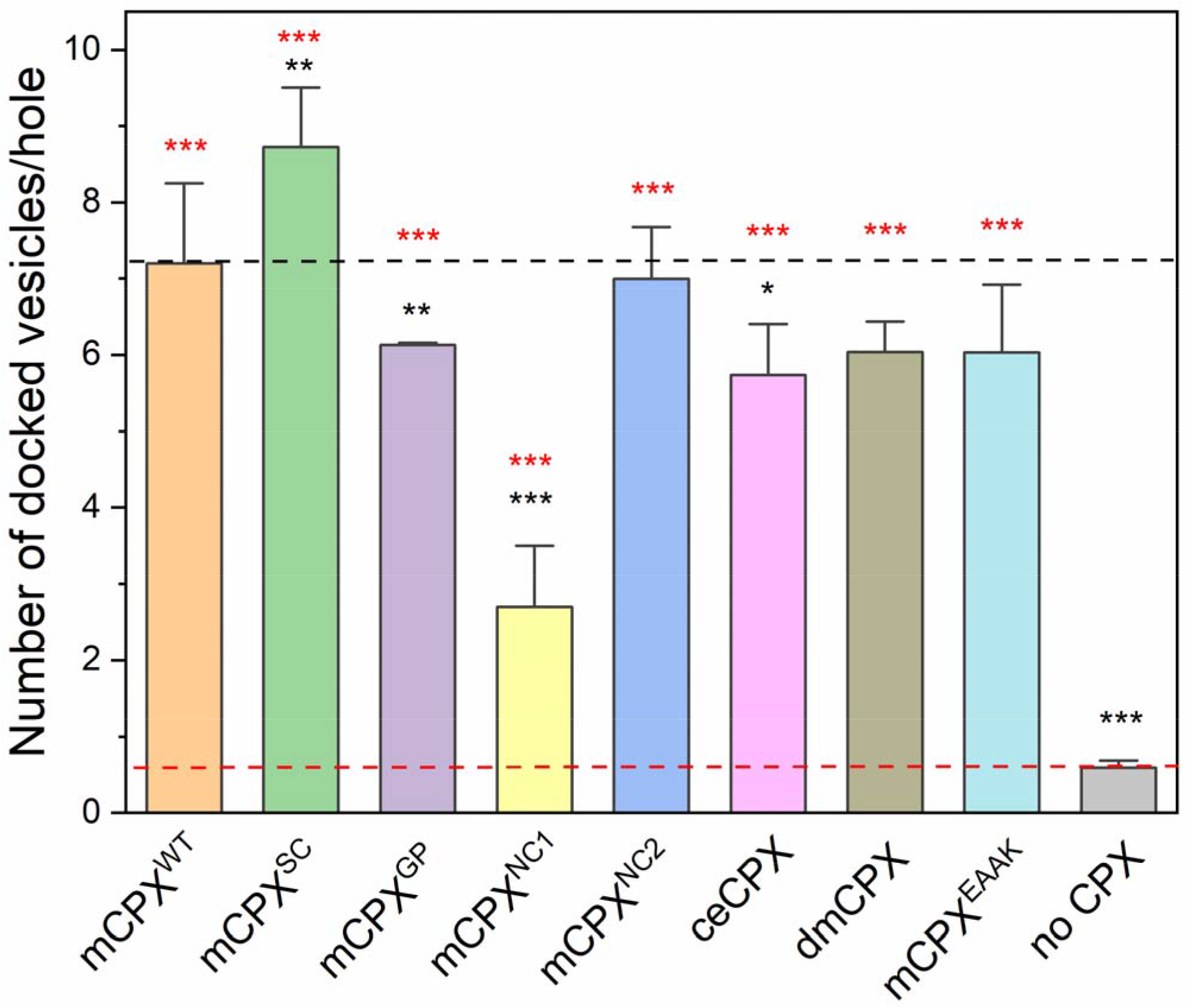
Docking analysis with Syt1-vSUVs show that CPX_acc_ does not contribute significantly to stimulatory effect on vesicle docking. With exception of the t-SNARE binding mutant (mCPX_NC1_), all other accessory helix mutants tested at the standard 2 μM concentration were able to significantly increase the number of vesicles docked. mCPX orthologs from Drosophila (dmCPX) and C. elegans (ceCPX) were also able to promote vesicle docking and to levels comparable to mCPX^WT^. Considering that CPX_cen_ is unaltered in mCPX mutants tested and shows high degree of sequence conservation in the orthologs, this data further supports the notion the CPX_cen_-SNARE interaction is the key to CPX ability to promote vesicle docking. In all experiments, VAMP2^4X^ mutant was used to enable unambiguous counting of stably docked vesicles. The average values and standard deviations from three independent experiments (with ~200 vesicles in total for each condition) are shown. *p<0.05, ** p<0.01 *** p<0.001 using the Student’s t-test.

**Figure 3: Supplement 3.**
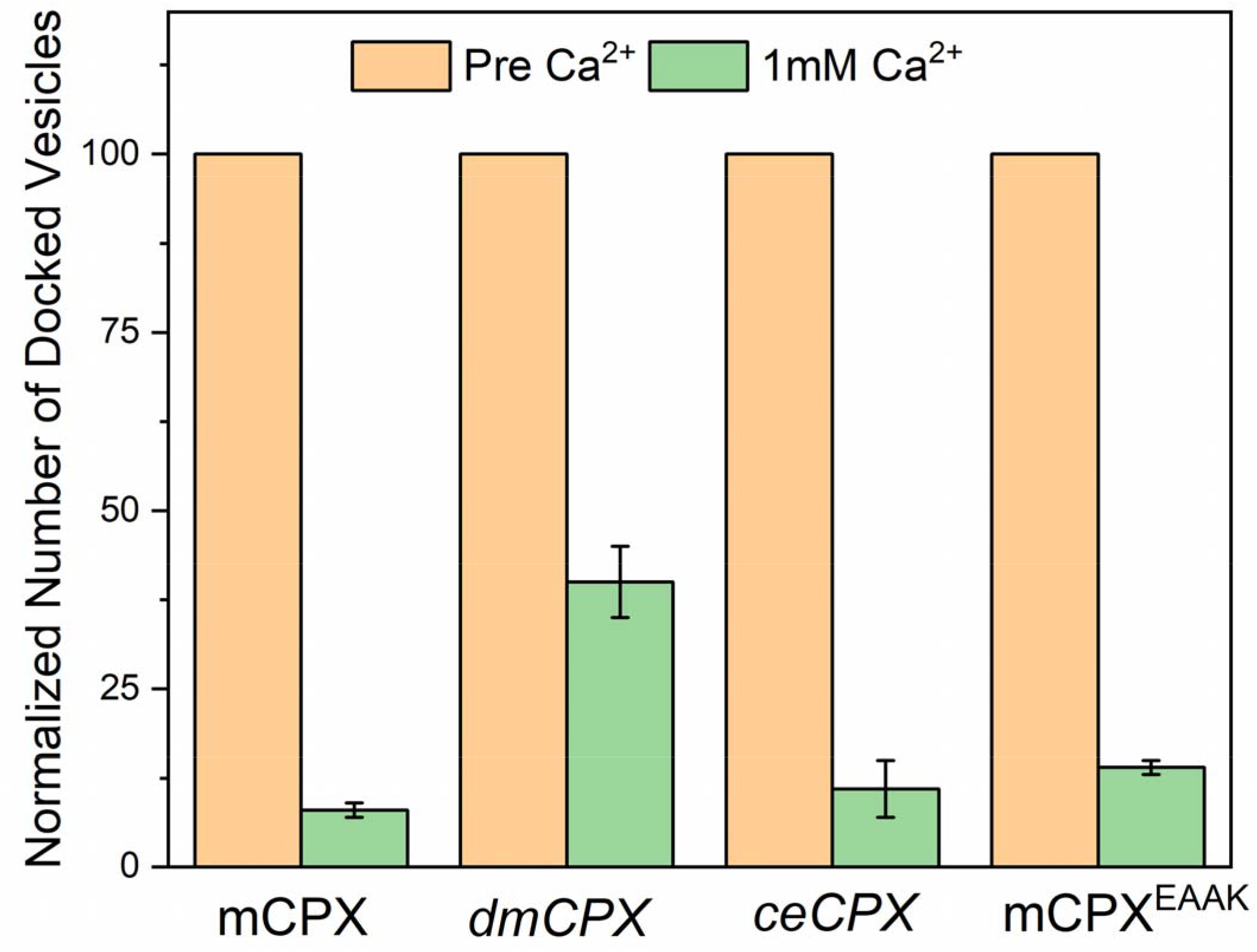
Syt1-vSUVs stably clamped at high concentration (20 μM) of Drosophila CPX (dmCPX), C. elegans CPX (ceCPX) and mCPX^EAAK^ remain Ca^2+^-sensitive and majority is triggered to fuse rapidly and synchronously following the addition of 1mM Ca^2+^. In these experiments, fusion was attested by sudden increase, followed by diffusion of ATTO647N-PE fluorescence introduced in the Syt1-vSUVs. Average and standard deviations from three independent experiments (minimum of 100 vesicles under each condition) are shown.

